# Discovering Novel Antimicrobial Peptides in Generative Adversarial Network

**DOI:** 10.1101/2021.11.22.469634

**Authors:** Tzu-Tang Lin, Li-Yen Yang, Ching-Tien Wang, Ga-Wen Lai, Chi-Fong Ko, Yang-Hsin Shih, Shu-Hwa Chen, Chung-Yen Lin

## Abstract

Due to the growing number of clinical antibiotic resistance cases in recent years, novel antimicrobial peptides (AMPs) can become ideal for next-generation antibiotics. This study trained a deep convolutional generative adversarial network (GAN) with known AMPs to generate novel AMP candidates. The quality of the GAN-designed peptides was evaluated *in silico*, and eight of them named GAN-pep 1∼8 were chosen to be synthesized for further experiments. Disk diffusion testing and minimum inhibitory concentration (MIC) determination were used to determine the antibacterial effects of the synthesized GAN-designed peptides. Seven out of the eight synthesized GAN-designed peptides showed antibacterial activities.

Additionally, GAN-pep 3 and GAN-pep 8 had a broad spectrum of antibacterial effects. Both of them were also effective against antibiotic-resistant bacteria strains such as methicillin-resistant *Staphylococcus aureus* (*S. aureus*) and carbapenem-resistant *Pseudomonas aeruginosa* (*P. aeruginosa*). GAN-pep 3, the most promising GAN-designed peptide candidate, had low MICs against all the tested bacteria.

## INTRODUCTION

The increasing number of clinical antibiotic resistance cases in the past decades raised the demand for new drug discovery of antibiotics (1, 2). AMPs are natural peptides that are less likely to cause drug resistance in bacteria (3, 4). However, it is usually time-consuming and costly to discover new AMPs through the traditional approaches by collecting these peptides from various organisms. Therefore, a deep learning model was proposed approach in this study for *in silico* AMP design to accelerate the AMP discovery process.

Artificial intelligence technologies and machine learning applications (AI/ML) are key to accelerating the drug development process (5, 6). For example, AI/ML can help initial drug selection for particular diseases. Zeng et al. used a knowledge graph embedding model to prioritize potential candidates for developing Coronavirus disease 2019 (COVID-19) therapy (7). Jiang et al. utilized a convolutional graph network to predict synergistic drug combinations against cancers (8). Also, AI/ML can be applied to predict various biological and chemical properties such as protein structure, molecular property, aqueous solubility, and minimum inhibitory concentration (MIC) (9-12). Moreover, AI/ML can be used to construct biomolecule classifiers to identify protein family, surfaceome, protein-protein interaction, human leukocyte antigen complex, and AMP (13-19). A deep neural network (DNN) named AI4AMP was recently proposed to predict the AMP activity (20).

In recent years, several researchers adopted *in silico* methods to support and accelerate AMP candidates finding. Some of the methods were directly based on computational algorithms (21, 22), and more and more studies utilized DNNs to generate peptides for AMP design. For example, Müller et al. trained a generative long short-term memory (LSTM) model to capture patterns of the AMP (23). Dean and Walper utilized variational autoencoder (VAE) to generate a latent space for creating new AMPs from known AMPs (24). Recently, the GAN, a neural network architecture composed of a generative model and a discriminative model, was used for DNA and protein design (25-29). Specifically, the deep convolutional GAN (DCGAN) was applied in various tasks of images generation (30).

The collected AMPS were first encoded by a protein-encoding method named PC6 proposed in the previous study, which turned each peptide into a matrix that considered the order of amino acids and the physicochemical properties of each amino acid (20). Then, a DCGAN-based model was proposed to generate AMPs (30). After training the model with these encoded AMPs, the generator could generate AMP candidates with random noise as its input. Then, these peptides were predicted through AI4AMP to evaluate the activity of the generated AMP candidates *in silico* before the experiment (20). Eight GAN-designed peptides named GAN-pep 1∼8 and a known AMP with high antimicrobial activity (AMP-pos) were used to test their activities against *Escherichia coli* (*E. coli*), *Staphylococcus aureus* (*S. aureus*), and *Pseudomonas aeruginosa* (*P. aeruginosa*) through the disk diffusion method and minimum inhibitory concentration (MIC) determination (31, 32). Moreover, to estimate the ability of the GAN-designed peptides against antibiotic-resistant bacteria, we further experimented with them on methicillin-resistant *S. aureus* and carbapenem-resistant *P. aeruginosa*.

## MATERIALS AND METHODS

### Collecting AMPs for training the model

The anti-bacterial AMPs from four AMP databases were collected (33-36). The AMPs with lengths shorter than ten amino acids or having uncommon amino acids, such as B, J, O, U, Z, or X, were excluded. According to the difficulty and the cost of synthesizing long peptides, only the AMPs shorter than 30 amino acids were selected. The number of collected AMPs was 3195.

### The architecture of the proposed GAN

The basic structure of GAN consists of a discriminator and a generator (25). The discriminator aims to discriminate between actual data and fake data generated by the generator. The generator, on the contrary, tries to generate fake data that can fool the discriminator. Meanwhile, the discriminator will be updated to maximize the discriminator score of the real data and minimize the score of the fake data. Oppositely, the generator is updated to maximize the discriminator score. The proposed GAN model for generating AMP was based on DCGAN, a convolutional network-based GAN (30). The kernel size, stride, and padding parameters in convolution-transpose layers and convolution layers were adjusted to fit the size of the data. The method proposed in WGAN-GP was used to avoid mode collapse (37).

The proposed generator consisted of five transposed convolution blocks. The first four building blocks were composed of a 2D-transposed convolution layer, a 2D-batch normalization layer, and an activation layer named Rectifier Linear Unit (ReLU). The last two transposed convolution blocks were composed of a 2D-transposed convolution layer and a Tanh activation layer. Five convolution blocks constructed the proposed discriminator by the first four building blocks of a 2D-convolutional layer and a leaky ReLU with a 2D-convolutional layer as the last block. The training data was first converted into vectors with shapes of (1, 30, 6), denoted as real PC6 matrix, by the method described in the next section. The generator took a noise vector with the shape of (100, 1, 1) and mapped it to the vector with the shape of (1, 30, 6), denoted as a fake PC6 matrix. The discriminator took in either the real PC6 matrix or the fake PC6 matrix and converted it into a vector with the shape of (1, 1, 1), which represented the discriminator score of the data. The proposed architectures of the generator and discriminator are shown in **Figure 1**. The K indicates the kernel size, and the S indicates the stride value.

**Figure 1.**
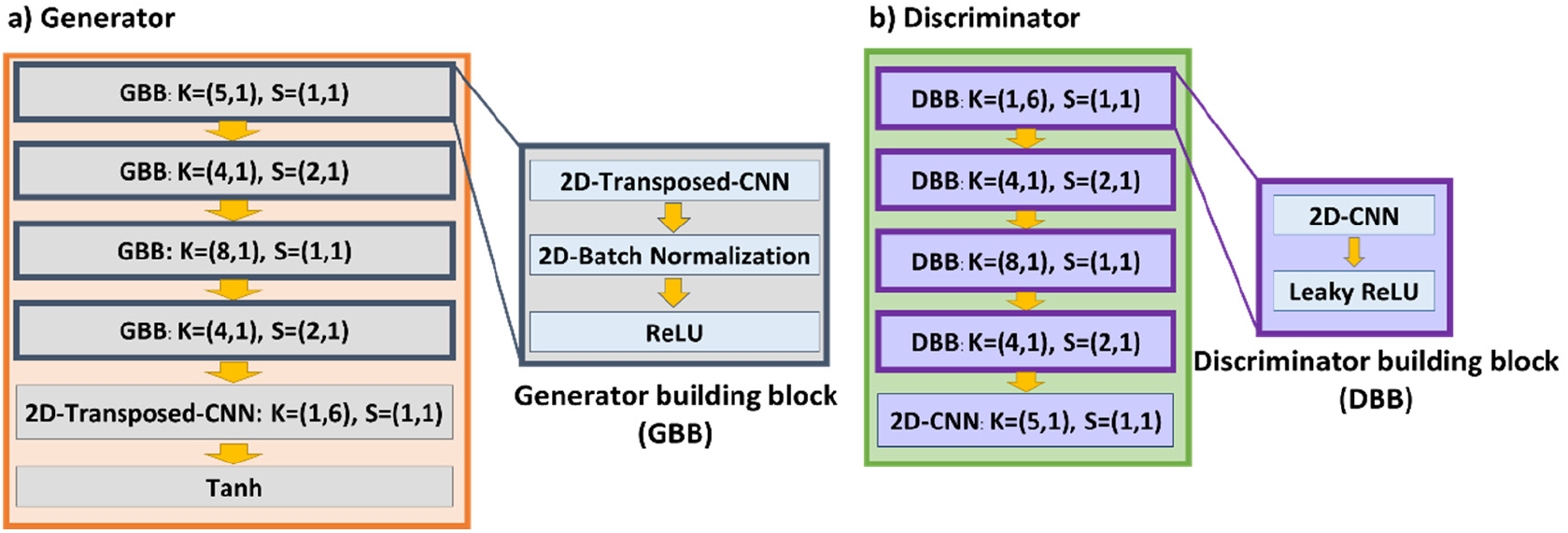
The proposed architecture of the generator and the discriminator.

### The mechanism of AMP production

For transforming peptides into numeric matrices, the PC6 protein-encoding method in the previous research was used to encode the peptides (20). The PC6 protein-encoding method required a PC6 table, where each amino acid corresponded to its six physicochemical properties values. This PC6 protein-encoding method transformed a peptide of length k into a matrix with the shape of (6, k). Six physicochemical property values in the PC6 table were scaled to a range of -1 to 1, respectively, to ensure every property had a balanced effect numerically in model training and to fit the tanh activation function in the last layer of the generator. For sequences shorter than 30 residues, they were padded with zero vector named “X” at the end to reach a consistent length of 30. Each AMP was then transformed into a real PC6 matrix with a shape of (1, 30, 6) by the scaled PC6 table. The real PC6 matrice was fed into the discriminator and produced discriminator scores. The fake PC6 matrices described in the previous paragraph were fed into the discriminator and produced the discriminator scores. The cosine similarity was used to convert the generated peptide from the fake PC6 matrix. The six generated physicochemical values of each row were converted into an amino acid with the highest similarity. If the six generated physicochemical values in the row were all zeros, the row would be converted into “X.” Any amino acid behind the first “X” and including itself would be discarded. **Figure 2** shows the overall workflow of training GAN to generate AMP.

**Figure 2.**
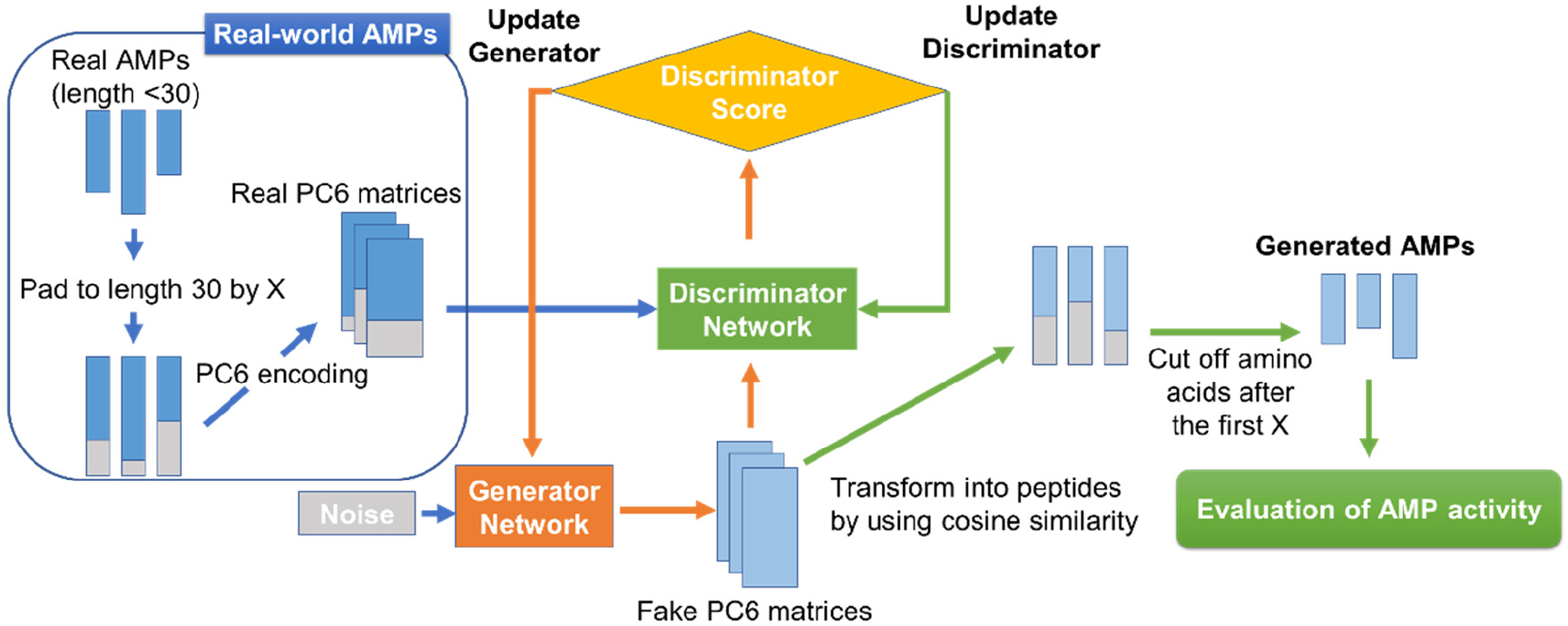
The overall workflow of the training GAN to generate AMP.

### The training process

Following WGAN-GP, the ratio of training steps between generator and discriminator was set to 1:5 (37). The batch size was set to 128. The Adam algorithm was applied as the optimizer for both models, with the learning rate being 1e-4, *β*_1_ being 0, and *β*_2_ being 0.9 (38). For every 5000 epochs, the 128 generator-designed sequences were evaluated. A fixed noise vector was used as the input for these generators. The outputs of these generators were transformed into peptides as described previously. After that, the identity between the generated peptide to the real AMP was evaluated by comparing the ratio of the same amino acid among the overlapped part. Each generated peptide was compared with every AMP in the dataset, producing 3195 identity scores. The identity score for the generated AMP was defined as the maximum identity it scored within the real AMP dataset. The number of the training process was 60,000 epochs. As shown in **Figure 3**, the identity score of the 128 test sequences produced by the current generators improved along with increased training steps, and it was stable around 50,000 epochs of training.

**Figure 3.**
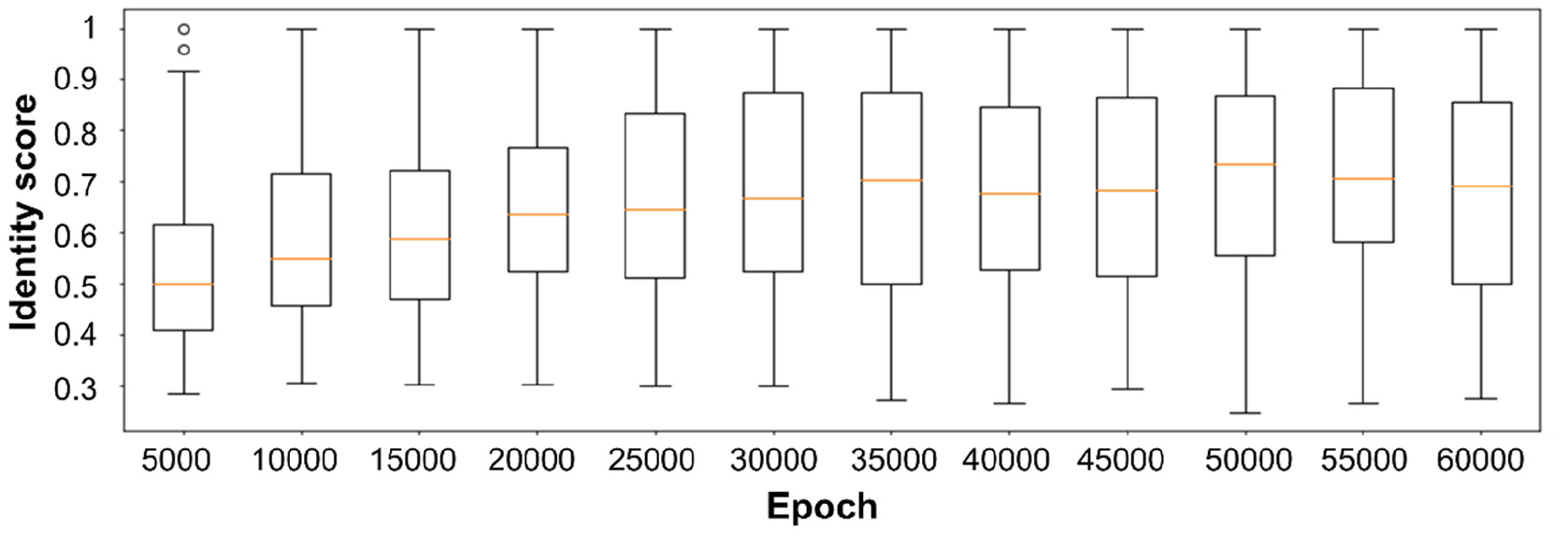
The boxplot of the maximum identity score distribution of generated peptides and real AMPs throughout the training process.

### Evaluation of GAN-designed sequences

To evaluate if the proposed GAN model had learned to generate peptides that had similar properties with real AMPs, the peptide properties of the GAN-designed peptides, the real AMPs, the randomly shuffled sequences, and the helical sequences were used to compare with real AMPs. The randomly shuffled sequences were peptides generated randomly with equal probabilities of all residues to ensure the proposed model did not merely generate random sequences. Since a large proportion of AMPs were composed of alpha helices, it would be interesting to see whether the model only learned the patterns of helices instead of the patterns to have antimicrobial properties. The generated peptides were compared with helical sequences generated by placing lysine or arginine every three or four amino acids. Both randomly shuffled sequences and helical sequences were generated with lengths of 10 to 30 by the modlAMP package (39). To perform the comparison to 3195 real AMPs, 3195 randomly shuffled sequences, 3195 helical sequences, and 3195 GAN-designed peptides were generated.

### GAN-designed sequences selection for experimental validation

By removing the duplicated peptides from 3195 GAN-designed peptides, 1970 GAN-designed peptides were left. Eight out of 1970 GAN-designed peptides were selected to test whether the produced sequences had antimicrobial activities by the following criteria. The GAN-designed peptides were kept only if eight physicochemical properties all fell in the range of the mean value plus or minus one standard deviation of those of the real antimicrobial peptides. The eight physicochemical properties were charge, charge density, isoelectric point, instability index, aromaticity, aliphatic index, Boman index, and hydrophobic ratio (40). These physicochemical properties were calculated by the modlamp package (39). After that, the remaining produced sequences were fed into AI4AMP (20), a CNN model that predicts the probability of a peptide with antimicrobial activity. The GAN-designed peptide was selected if its probability of having antimicrobial activities was greater than 0.98. Based on the identity scores of the 1970 GAN-designed peptides, they were classified into three categories. The very similar sequences were sequences with identity scores from 80% to 98%. The moderately similar sequences were sequences with identity scores from 40% to 60%. The dissimilar sequences were sequences with identity scores smaller than 20%. To test if sequences that were not so similar to the real AMPs were still able to possess antibacterial properties, 21 sequences from the very similar sequence category were selected, and 13 sequences from the moderately similar sequence category were selected. No sequences were selected from the dissimilar sequence category. Then, four sequences from the very similar sequence category (GAN-pep 1∼4) were chosen, and four sequences from the moderately similar sequence category (GAN-pep 5∼8) were chosen. These eight peptides were chosen to be synthesized for further antimicrobial experiments.

### Strains and Reagents

The bacterial strains used for antimicrobial activity assays include *E. coli* (SG13009), the clinical isolates of methicillin-susceptible *S. aureus* (S01-10-0202), methicillin-resistant *S. aureus* (N07-10-0043), carbapenem-susceptible *P. aeruginosa* (S07-10-0059), and carbapenem-resistant *P. aeruginosa* (M06-06-0213), obtained from Dr. Ying-Lien Chen, Department of Plant Pathology and Microbiology, National Taiwan University. All strains were grown aerobically on an orbital shaker (150 rpm) at 37 °C in Luria-Bertani (LB) broth (BD Difco, US) overnight. Different microbiological assays were performed to test their antimicrobial activity with the GAN-designed peptides (GAN-pep 1∼8) and the positive control peptide (AMP-pos). Experiments were performed in triplicate.

### Antimicrobial assays

The antibacterial potential of the GAN-designed peptides was evaluated using disk diffusion assay. The bacteria were grown in LB broth at 37°C with agitation. The strain growth was measured turbid metrically at OD_600_, and at least three separate experiments were conducted for each test organism. Briefly, nutrient agar was prepared by mixing agar, sodium chloride, yeast extract, and peptone in distilled water (pH 7.2). Subsequently, bacterial suspension (100 μL, 1 × 10^8^ CFU/mL) was added and spread on LB agar. Then, the sterilized filter disks (with diameter circles of 6 mm) were placed on the agar surface filled with 40 μL of peptide samples. The petri dish was incubated overnight at 37 °C to observe the inhibitory area.

MIC assays were conducted to determine the antibacterial spectrum of these peptides. The MIC was determined as the lowest peptide concentration inhibited bacterial growth after overnight incubation at 37 °C. Microbial strains were cultured in LB medium, and mid-logarithmic-phase organisms were used in antibacterial assays. All bacteria were inoculated in LB medium (approximately 10^5^ CFU/mL), and MIC assays were performed with different concentrations of each peptide. All activity measurements were conducted at least three times.

## RESULTS AND DISCUSSION

### Evaluating GAN-designed peptides *in silico*

**Figure 4**. shows the amino acid distribution of four groups of peptides. It showed that the amino acid composition of real AMPs and the GAN-designed peptides had a very similar pattern compared to the other two groups, which suggests that the GAN model can capture the pattern in terms of the sequence composition. This result indicates that the model is neither merely generating random sequences nor learning the patterns for alpha helix. **Figure 5** shows the violin plots of eight physicochemical properties of the four groups of peptides. The “AMP” indicates the real AMPs, “GAN” indicates the GAN-designed peptides, “Random” indicates the randomly shuffled sequences, and “Helical” indicates the helical sequences. The eight physicochemical properties used to evaluate the peptides were aliphatic index, aromaticity, Boman index, charge density, charge, hydrophobic ratio, instability index, and isoelectric point (40). The distribution pattern of the GAN-designed peptides resembled that of the real AMPs. Such a pattern suggests that the GAN model is possible to produce peptides with physicochemical properties. The eight physicochemical features were then reduced into three dimensions by t-distributed stochastic neighbor embedding (t-SNE) and were visualized by Matplotlib (41, 42). **Figure 6** shows the t-SNE plot for the four groups of peptides. The “AMP” indicates real AMPs, “GAN” indicates GAN-designed peptides, “Random” indicates randomly shuffled sequences, and “Helical” indicates helical sequences. It showed that the real AMPs and the GAN-designed peptides were clustered closely, distinct from the two other groups in embedded space, showing that the GAN-designed peptides possessed very similar properties with the real AMPs.

**Figure 4.**
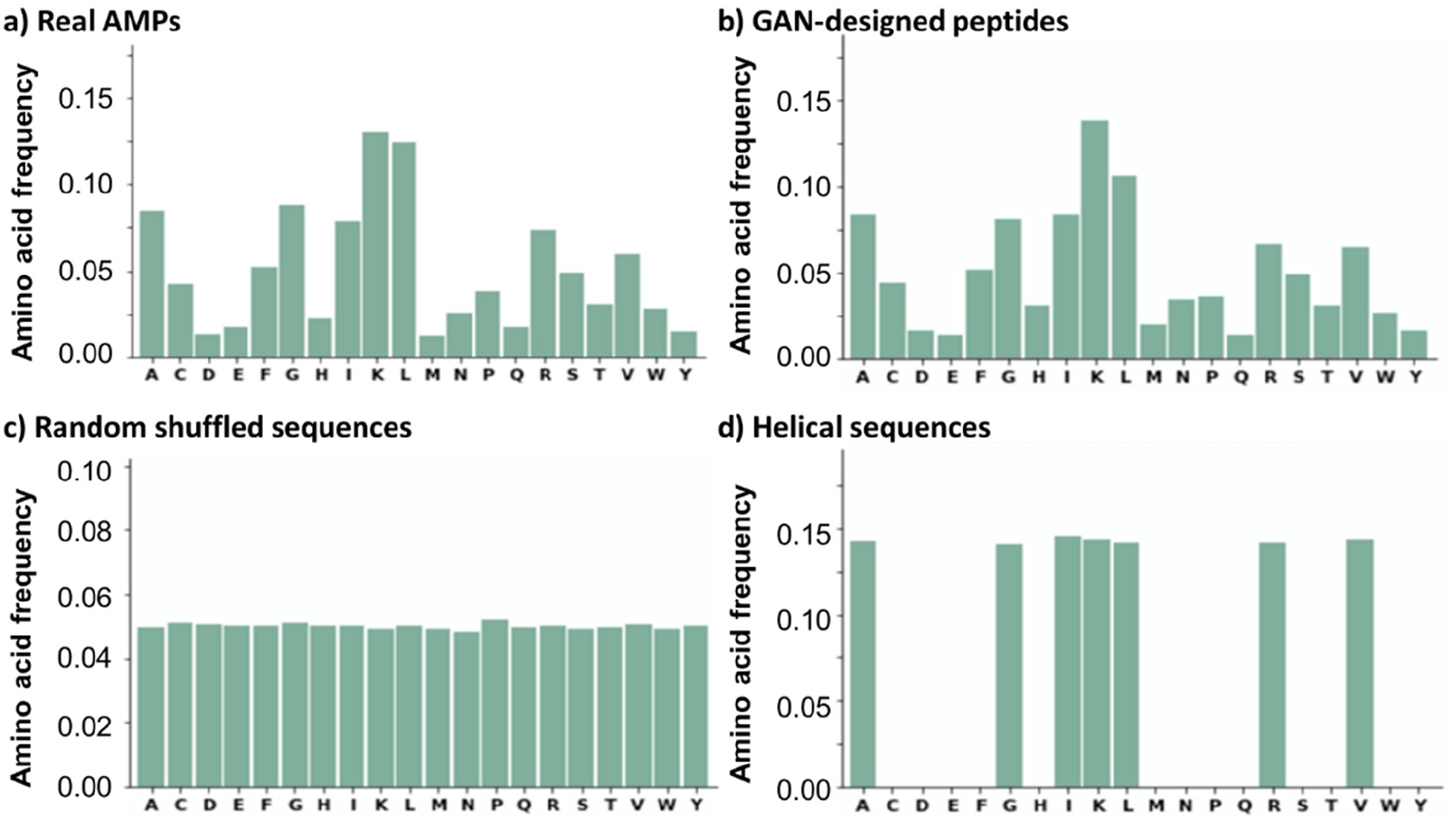
Bar plots of amino acid distribution in four groups of peptides.

**Figure 5.**
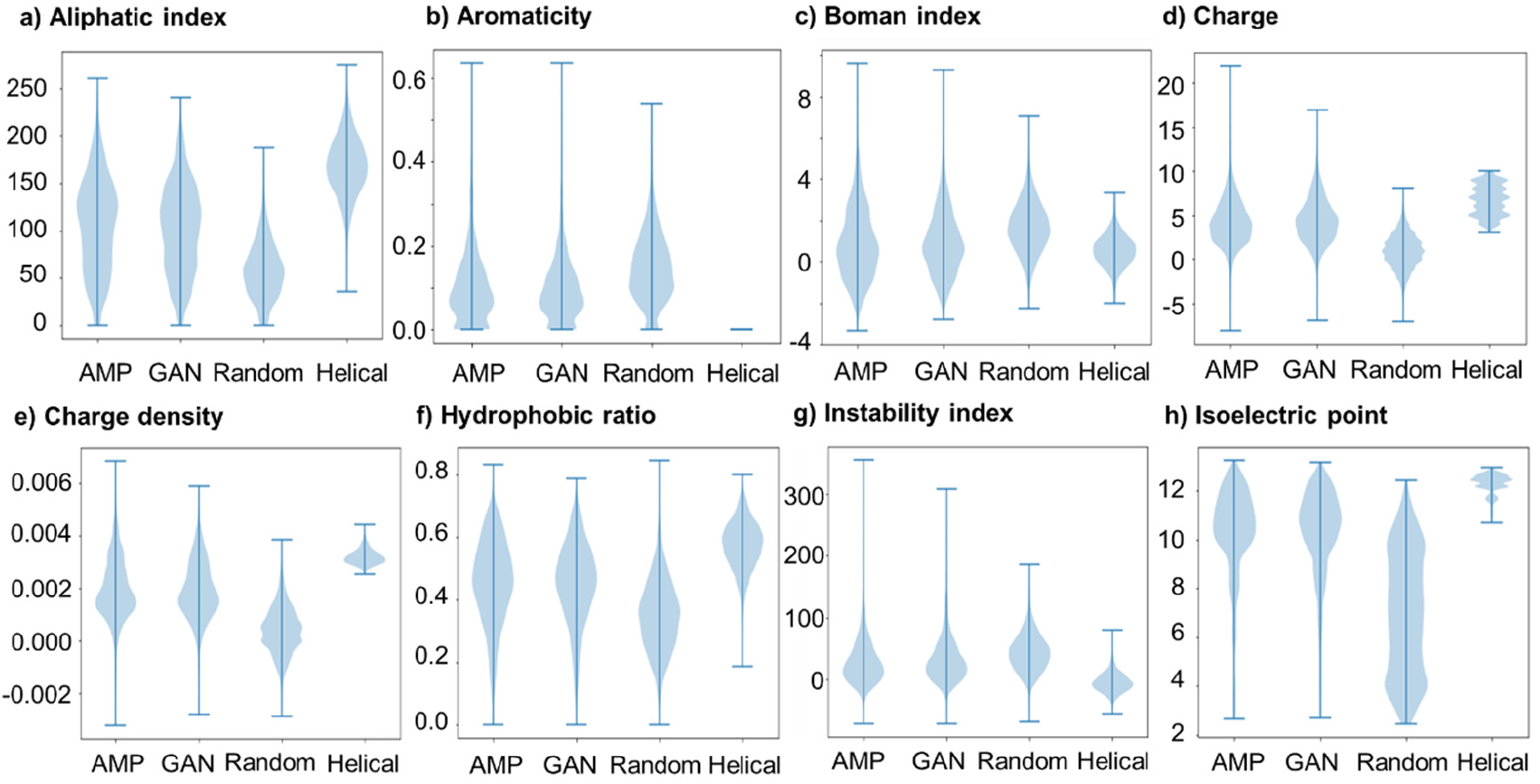
Violin plots of physicochemical properties in four groups of peptides.

**Figure 6.**
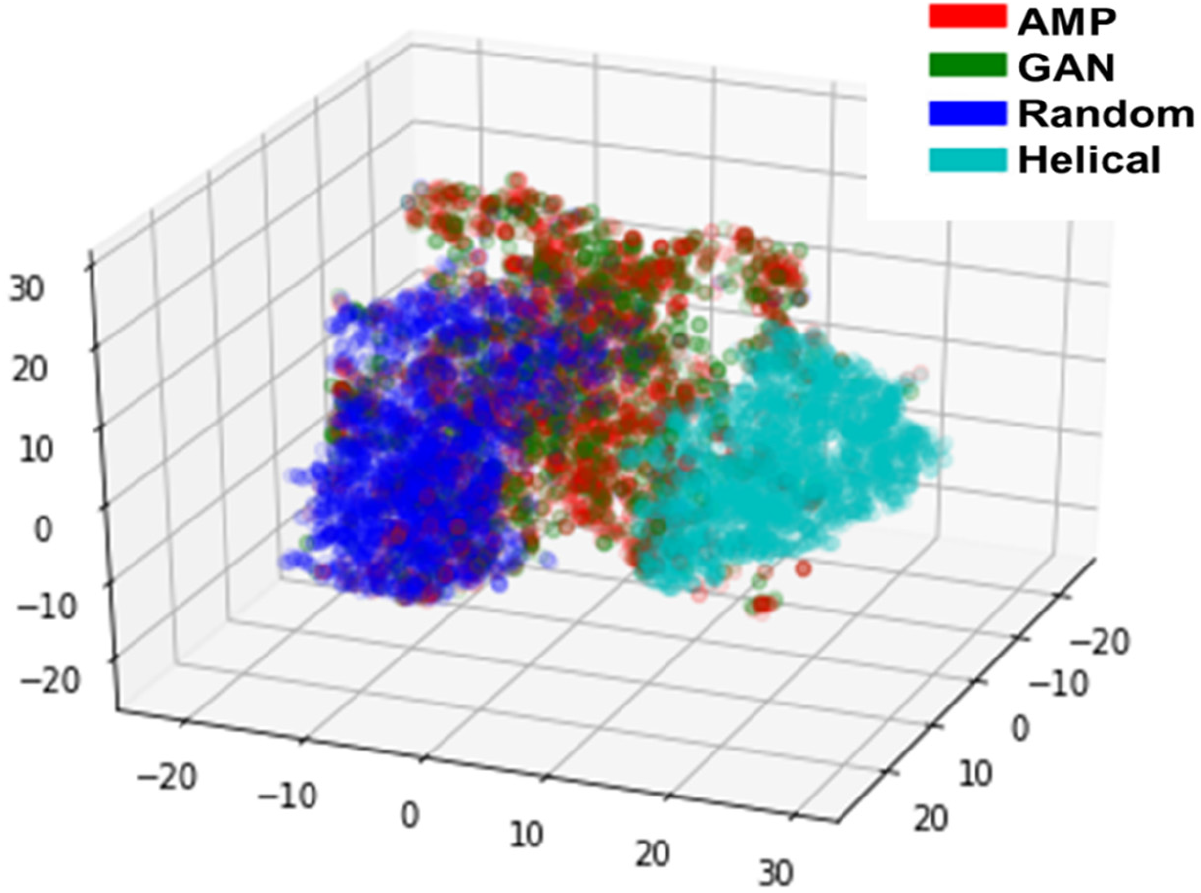
The t-SNE plot of four groups of peptides.

### Evaluating GAN-designed peptides *in vitro*

Various concentrations (7.8125 to 500 μg/mL) of GAN-designed peptides (GAN-pep 1∼8), a known AMP (AMP-pos) as the positive control, and bovine serum albumin (BSA) as negative control were prepared for disk diffusion assay. The results for the disk diffusion susceptibility test of GAN-designed peptides, and the positive control peptide, and the negative control peptide at different concentrations against several bacteria, including *E. coli*, the clinical isolates of methicillin-susceptible *S. aureus*, methicillin-resistant *S. aureus*, carbapenem-susceptible *P. aeruginosa*, and carbapenem-resistant *P. aeruginosa*, are shown in **Supplementary Figure 1 to 5**. As presented in **Supplementary Figure 1**, for AMP-pos and some of the GAN-designed peptides, such as GAN-pep 2, 3, 4, 5, 7, and 8, at least one concentration of the peptides inhibited the tested Gram-negative bacterium *E. coli*. As presented in **Supplementary Figure 2**, for AMP-pos and some of the GAN-designed peptides, such as GAN-pep 3, 4, 6, and 8, at least one concentration inhibited the tested Gram-positive bacterium methicillin-susceptible *S. aureus*. As presented in **Supplementary Figure 3**, for AMP-pos and some of the GAN-designed peptides, such as GAN-pep 3, 6, and 8, at least one concentration of the peptides inhibited the tested Gram-positive bacterium methicillin-resistant *S. aureus*. As presented in **Supplementary Figure 4**, only the GAN-designed peptides, such as GAN-pep 2, 3, 4, and 8, could inhibit the tested Gram-negative bacterium carbapenem-susceptible *P. aeruginosa* at one or more concentrations. As presented in **Supplementary Figure 5**, only the GAN-designed peptides, such as GAN-pep 2, 3, and 8, could inhibit the tested Gram-negative bacterium carbapenem-resistant *P. aeruginosa* at one or more concentrations. Overall, GAN-pep 3 and GAN-pep 8 had the broadest antibacterial effects against all tested bacteria.

The MIC of each peptide for a selection of microorganisms is shown in **Table 1**. The AMP-pos and GAN-designed peptides, such as GAN-pep 2, 3, 4, 5, 7, and 8, had MIC ranging from 0.7 to 22.5 μg/mL against the tested Gram-negative bacterium *E. coli*. AMP-pos showed the highest antibacterial activity against *E. coli*. with MIC of 0.7 μg/mL. The GAN-designed peptides, GAN-pep 3 and 8, had MIC ranging from 6 to 15 μg/mL against the tested Gram-positive bacterium methicillin-susceptible *S. aureus*. The GAN-designed peptides, GAN-pep 3 and 8, had 45 μg/mL MIC against the tested Gram-positive bacterium methicillin-resistant *S. aureus*. The GAN-designed peptides, GAN-pep 2, 3, and 4, had MIC ranging from 3 to 50 μg/mL against the tested Gram-negative bacterium carbapenem-susceptible *P. aeruginosa*. The GAN-designed peptides, GAN-pep 2, 3, and 4, had MIC ranging from 3 to 35 μg/mL against the tested Gram-negative bacterium carbapenem-resistant *P. aeruginosa*.

**Table 1.** Antibacterial activity of GAN-designed peptides and two positive peptides.

| name | MIC (μg/mL) |  |  |  |  |
| --- | --- | --- | --- | --- | --- |
|  | *E. coli*<br>(SG13009) | methicillin-susceptible<br>*S. aureus*<br>(S01-10-0202) | methicillin-resistant<br>*S. aureus*<br>(N07-10-0043) | carbapenem-susceptible<br>*P. aeruginosa*<br>(S07-10-0059) | carbapenem-resistant<br>*P. aeruginosa*<br>(M06-06-0213) |
| AMP-pos | 0.7 | >50 | >50 | >50 | >50 |
| GAN-pep 1 | >50 | >50 | >50 | >50 | >50 |
| GAN-pep 2 | 2 | >50 | >50 | 50 | 5 |
| GAN-pep 3 | 2 | 6 | 45 | 3 | 3 |
| GAN-pep 4 | 2 | >50 | >50 | 50 | 35 |
| GAN-pep 5 | 22.5 | >50 | >50 | >50 | >50 |
| GAN-pep 6 | >50 | >50 | >50 | >50 | >50 |
| GAN-pep 7 | >50 | >50 | >50 | >50 | >50 |
| GAN-pep 8 | 15 | 15 | 45 | >50 | >50 |

Overall, seven out of eight GAN-designed peptides showed antimicrobial activity against at least one strain of the bacteria. This strategy demonstrates that the GAN model can design novel sequence patterns with antimicrobial activity. Among the GAN-designed peptides, GAN-pep 3 and GAN-pep 8 showed broad and practical antibacterial activities. GAN-pep 3 and GAN-pep 8 had the inhibition effect against both gram-negative and gram-positive bacteria. In addition to that, they were able to inhibit bacteria strains that have developed antibiotic resistance.

## CONCLUSION AND FUTURE WORK

In this study, a new AMPs design method was proposed to support AMP discovery. The anti-bacterial AMPs were encoded through the PC6 protein-encoding method and then be used to train the proposed GAN model using a modified DCGAN architecture based on WGAN-GP (30, 37). The trained generator then generated the AMP candidates. These AMP candidates were evaluated by comparing peptide amino acid distribution and physicochemical property between four types of peptide groups. In addition, a deep learning model named AI4AMP was used to predict the AMP activity of the GAN-designed peptides (20). The eight GAN-designed peptides (GAN-pep 1∼8) predicted to have antimicrobial activities with probabilities greater than 0.98 were synthesized. Finally, the AMP activities of GAN-pep 1∼8 were examined by using disk diffusion testing and MIC determination. Seven of the eight synthesized GAN-designed peptides showed antibacterial activities, which showed that the proposed GAN model could design AMPs with antibacterial effects. Among them, GAN-pep 3 and GAN-pep 8 possessed a broad spectrum of antibacterial effects. They were also effective against antibiotic-resistant bacteria strains such as methicillin-resistant *S. aureus* and carbapenem-resistant *P. aeruginosa*. GAN-pep 3, the most promising AMP candidates, had even lower MICs against *S. aureus* and *P. aeruginosa* than the positive control AMP.

In the hope of developing the GAN-designed peptides into potential drugs, further experiments should be done. Since hemolysis is one of the significant effects that cause safety concerns and hinders AMPs from processing through later phases of drug development, experiments for the hemolysis effect of those GAN-designed peptides should be tested. The proposed model could generate various kinds of short peptides. It might be used to design peptides with antiviral and antifungal effects.

## ACKNOWLEDGMENTS

The authors thank the Ministry of Science and Technology (MOST), Taiwan, and Academia Sinica, Taiwan, for financially supporting this study and publication through 108-2314-B-001-002, 108-2321-B-038-003, and Grand Challenge Seed Program, respectively. Conflict of Interest: none declared.

## CODE AVAILABILITY

The code for training the GAN model for AMP design is publicly available at: https://github.com/lsbnb/amp_gan.

## REFERENCE

1. Baker SJ, Payne DJ, Rappuoli R, De Gregorio E. 2018. Technologies to address antimicrobial resistance. Proc Natl Acad Sci U S A 115:12887–12895.

2. Aslam B, Wang W, Arshad MI, Khurshid M, Muzammil S, Rasool MH, Nisar MA, Alvi RF, Aslam MA, Qamar MU, Salamat MKF, Baloch Z. 2018. Antibiotic resistance: a rundown of a global crisis. Infection and drug resistance 11:1645–1658.

3. Spohn R, Daruka L, Lázár V, Martins A, Vidovics F, Grézal G, Méhi O, Kintses B, Számel M, Jangir PK, Csörgő B, Györkei Á, Bódi Z, Faragó A, Bodai L, Földesi I, Kata D, Maróti G, Pap B, Wirth R, Papp B, Pál C. 2019. Integrated evolutionary analysis reveals antimicrobial peptides with limited resistance. Nature Communications 10:4538.

4. Galdiero E, Lombardi L, Falanga A, Libralato G, Guida M, Carotenuto R. 2019. Biofilms: novel strategies based on antimicrobial peptides. Pharmaceutics 11:322.

5. Levin JM, Oprea TI, Davidovich S, Clozel T, Overington JP, Vanhaelen Q, Cantor CR, Bischof E, Zhavoronkov A. 2020. Artificial intelligence, drug repurposing and peer review. Nature Biotechnology 38:1127–1131.

6. Réda C, Kaufmann E, Delahaye-Duriez A. 2020. Machine learning applications in drug development. Computational and Structural Biotechnology Journal 18:241–252.

7. Zeng X, Song X, Ma T, Pan X, Zhou Y, Hou Y, Zhang Z, Li K, Karypis G, Cheng F. 2020. Repurpose open data to discover therapeutics for COVID-19 using deep learning. Journal of proteome research 19:4624–4636.

8. Jiang P, Huang S, Fu Z, Sun Z, Lakowski TM, Hu P. 2020. Deep graph embedding for prioritizing synergistic anticancer drug combinations. Computational and Structural Biotechnology Journal 18:427–438.

9. Senior AW, Evans R, Jumper J, Kirkpatrick J, Sifre L, Green T, Qin C, Žídek A, Nelson AW, Bridgland A. 2020. Improved protein structure prediction using potentials from deep learning. Nature 577:706–710.

10. Wang S, Guo Y, Wang Y, Sun H, Huang J. SMILES-BERT: large scale unsupervised pre-training for molecular property prediction, p 429–436. In (ed),

11. Chen J-H, Tseng YJ. 2020. Different molecular enumeration influences in deep learning: an example using aqueous solubility. Briefings in Bioinformatics.

12. Witten J, Witten Z. 2019. Deep learning regression model for antimicrobial peptide design. BioRxiv:692681.

13. Seo S, Oh M, Park Y, Kim S. 2018. DeepFam: deep learning based alignment-free method for protein family modeling and prediction. Bioinformatics 34:i254–i262.

14. Bausch-Fluck D, Goldmann U, Müller S, van Oostrum M, Müller M, Schubert OT, Wollscheid B. 2018. The in silico human surfaceome. Proceedings of the National Academy of Sciences 115:E10988–E10997.

15. Sun T, Zhou B, Lai L, Pei J. 2017. Sequence-based prediction of protein protein interaction using a deep-learning algorithm. BMC Bioinformatics 18:277.

16. Vang YS, Xie X. 2017. HLA class I binding prediction via convolutional neural networks. Bioinformatics 33:2658–2665.

17. Bhadra P, Yan J, Li J, Fong S, Siu SWI. 2018. AmPEP: Sequence-based prediction of antimicrobial peptides using distribution patterns of amino acid properties and random forest. Scientific Reports 8:1697.

18. Meher PK, Sahu TK, Saini V, Rao AR. 2017. Predicting antimicrobial peptides with improved accuracy by incorporating the compositional, physico-chemical and structural features into Chou’s general PseAAC. Scientific Reports 7:42362.

19. Veltri D, Kamath U, Shehu A. 2018. Deep learning improves antimicrobial peptide recognition. Bioinformatics 34:2740–2747.

20. Lin T-T, Yang L-Y, Lu I-H, Cheng W-C, Hsu Z-R, Chen S-H, Lin C-Y. 2020. AI4AMP: Sequence-based antimicrobial peptides predictor using physicochemical properties-based encoding method and deep learning. bioRxiv.

21. Porto WF, Irazazabal L, Alves ES, Ribeiro SM, Matos CO, Pires ÁS, Fensterseifer IC, Miranda VJ, Haney EF, Humblot V. 2018. In silico optimization of a guava antimicrobial peptide enables combinatorial exploration for peptide design. Nature communications 9:1–12.

22. Porto W, Fensterseifer I, Ribeiro S, Franco O. 2018. Joker: An algorithm to insert patterns into sequences for designing antimicrobial peptides. Biochimica et Biophysica Acta (BBA) - General Subjects 1862.

23. Müller AT, Hiss JA, Schneider G. 2018. Recurrent neural network model for constructive peptide design. Journal of chemical information and modeling 58:472–479.

24. Dean SN, Walper SA. 2020. Variational Autoencoder for Generation of Antimicrobial Peptides. ACS omega 5:20746–20754.

25. Goodfellow IJ, Pouget-Abadie J, Mirza M, Xu B, Warde-Farley D, Ozair S, Courville A, Bengio Y. 2014. Generative adversarial nets, abstr Proceedings of the 27th International Conference on Neural Information Processing Systems - Volume 2, Montreal, Canada, MIT Press,

26. Anand N, Huang P. 2018. Generative modeling for protein structures. Advances in Neural Information Processing Systems 31:7494–7505.

27. Rossetto AM, Zhou W. GANDALF: A Prototype of a GAN-based Peptide Design Method, p 61–66. In (ed),

28. Killoran N, Lee LJ, Delong A, Duvenaud D, Frey BJ. 2017. Generating and designing DNA with deep generative models. arXiv preprint 171206148.

29. Gupta A, Zou J. 2018. Feedback GAN (FBGAN) for DNA: a novel feedback-loop architecture for optimizing protein functions. arXiv preprint 180401694.

30. Radford A, Metz L, Chintala S. 2015. Unsupervised representation learning with deep convolutional generative adversarial networks. arXiv preprint 151106434.

31. Zhang L, Scott MG, Yan H, Mayer LD, Hancock RE. 2000. Interaction of polyphemusin I and structural analogs with bacterial membranes, lipopolysaccharide, and lipid monolayers. Biochemistry 39:14504–14514.

32. Andrews JM. 2001. Determination of minimum inhibitory concentrations. Journal of antimicrobial Chemotherapy 48:5–16.

33. Li X, Wang Z, Wang G. 2015. APD3: the antimicrobial peptide database as a tool for research and education. Nucleic Acids Research 44:D1087–D1093.

34. Zhao X, Wu H, Lu H, Li G, Huang Q. 2013. LAMP: A Database Linking Antimicrobial Peptides. PLOS ONE 8:e66557.

35. Waghu FH, Barai RS, Gurung P, Idicula-Thomas S. 2016. CAMPR3: a database on sequences, structures and signatures of antimicrobial peptides. Nucleic Acids Research 44:D1094–D1097.

36. Kang X, Dong F, Shi C, Liu S, Sun J, Chen J, Li H, Xu H, Lao X, Zheng H. 2019. DRAMP 2.0, an updated data repository of antimicrobial peptides. Scientific Data 6:148.

37. Gulrajani I, Ahmed F, Arjovsky M, Dumoulin V, Courville AC. 2017. Improved training of wasserstein gans. Advances in neural information processing systems 30:5767–5777.

38. Kingma DP, Ba JL. 2015. Adam: A Method for Stochastic Optimization, abstr International Conference on Learning Representations, 1/1/2015.

39. Müller AT, Gabernet G, Hiss JA, Schneider G. 2017. modlAMP: Python for antimicrobial peptides. Bioinformatics 33:2753–2755.

40. Boman HG. 2003. Antibacterial peptides: basic facts and emerging concepts. Journal of Internal Medicine 254:197–215.

41. Maaten Lvd, Hinton G. 2008. Visualizing data using t-SNE. Journal of machine learning research 9:2579–2605.

42. Hunter JD. 2007. Matplotlib: A 2D graphics environment. Computing in science & engineering 9:90–95.

