## Supplements for "Discovering Novel Antimicrobial Peptides in Generative Adversarial Network"

### Table of Contents of Supplementary Figures

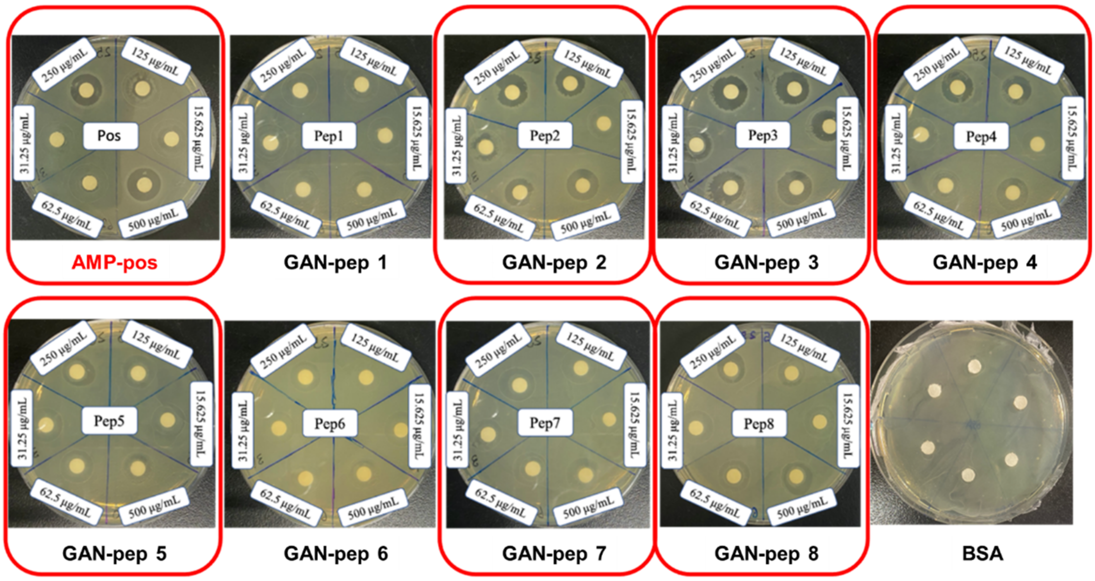

**Supplementary Figure 1.** Growth inhibition test against *E. coli* with peptides at different concentrations. Peptides are highlighted with red rectangles if inhibition zones occur around the disks.

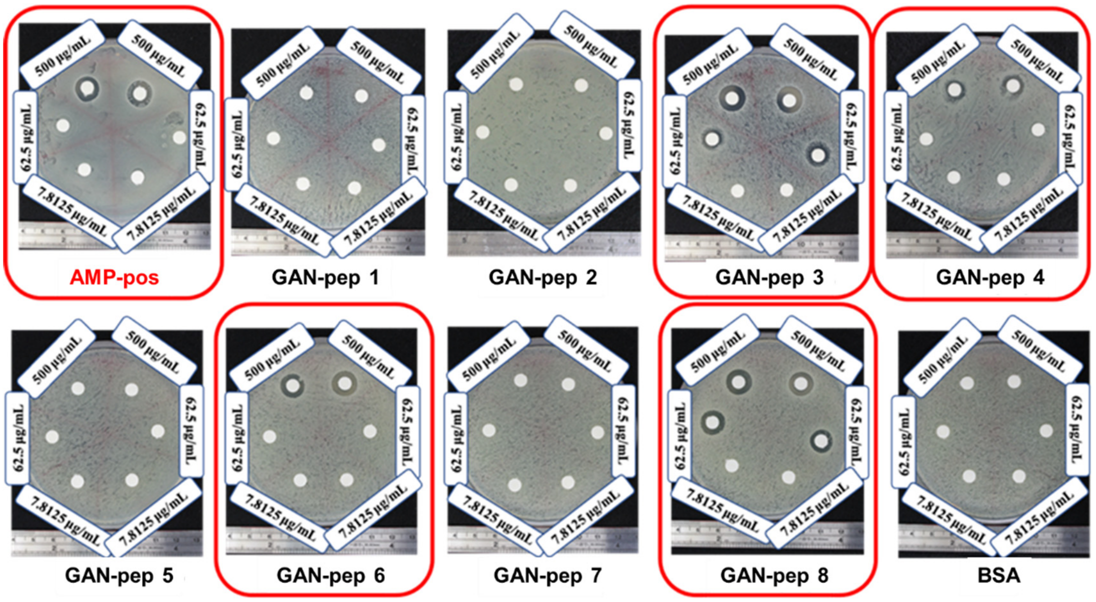

**Supplementary Figure 2.** Growth inhibition test against methicillin-susceptible *S. aureus* with peptides at different concentrations. Peptides are highlighted with red rectangles if inhibition zones occur around the disks.

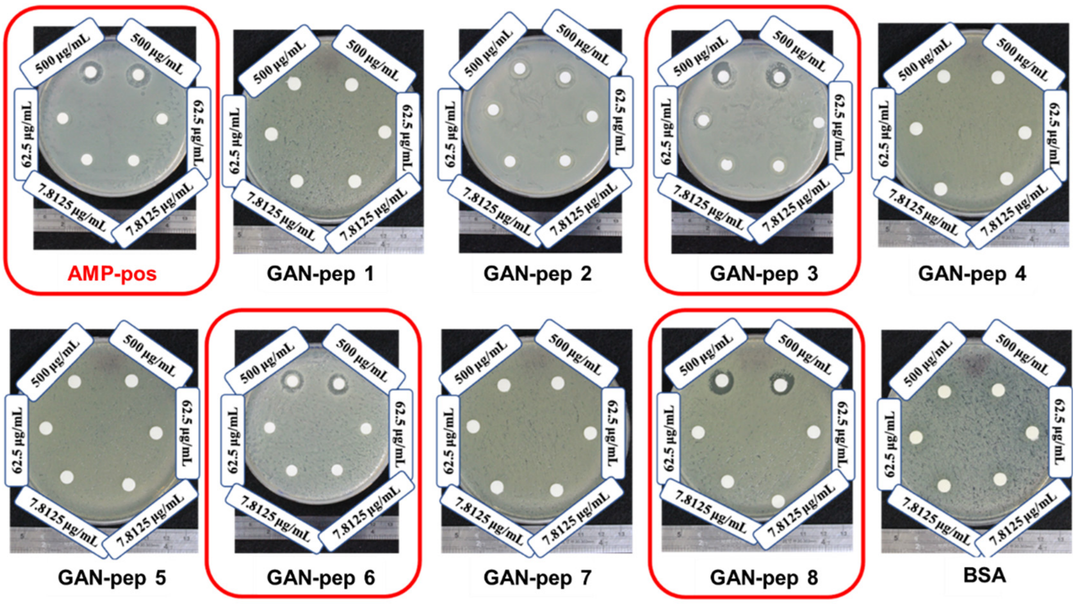

**Supplementary Figure 3.** Growth inhibition test against methicillin-resistant *S. aureus* with peptides at different concentrations. Peptides are highlighted with red rectangles if inhibition zones occur around the disks.

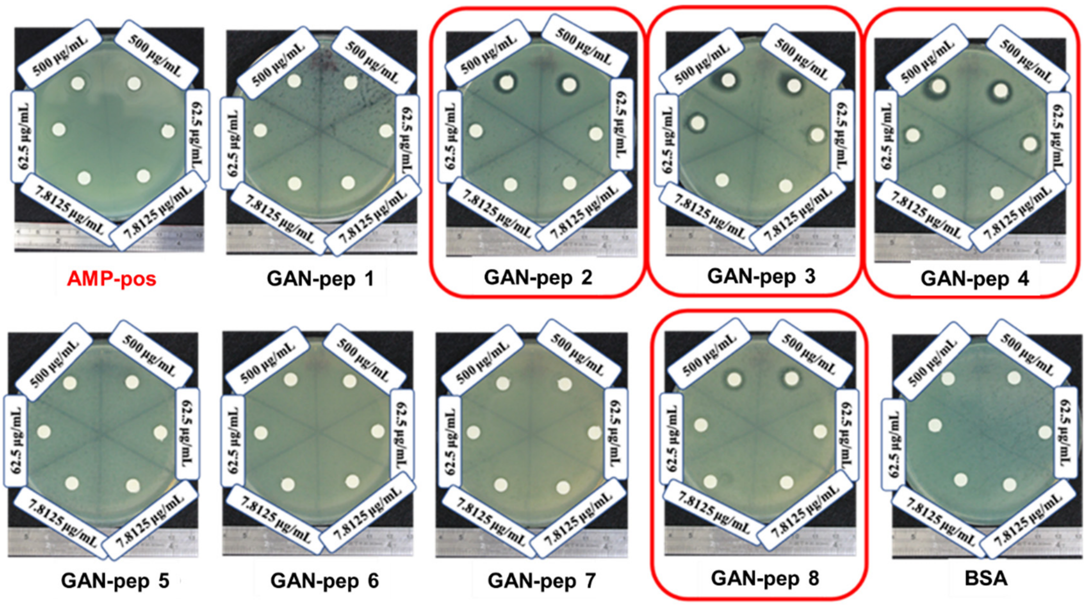

**Supplementary Figure 4.** Growth inhibition test against carbapenem-susceptible *P. aeruginosa* with peptides at different concentrations. Peptides are highlighted with red rectangles if inhibition zones occur around the disks.

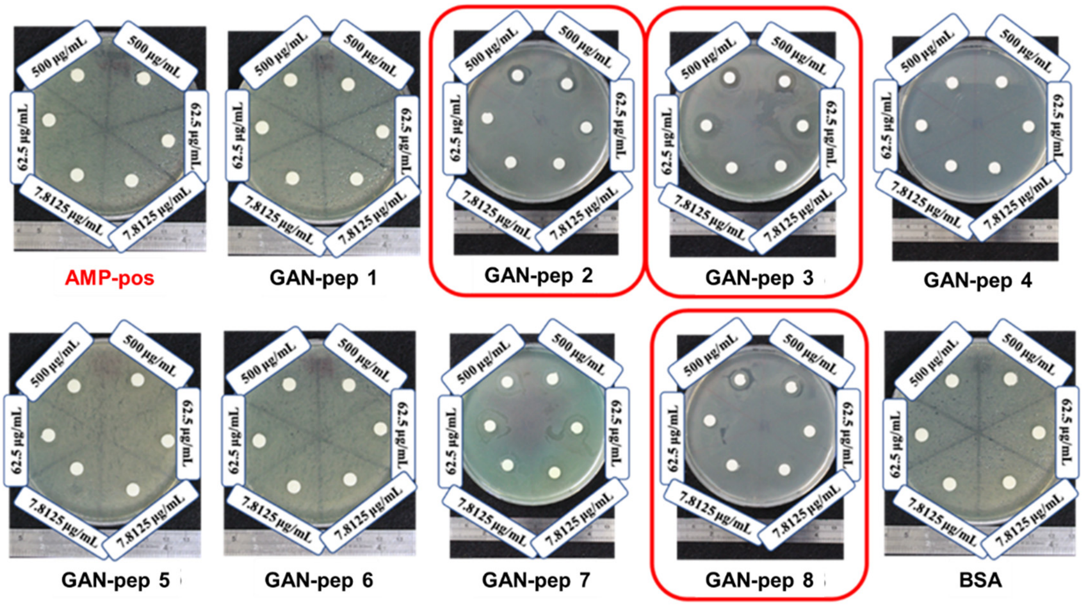

**Supplementary Figure 5.** Growth inhibition test against carbapenem-resistant *P. aeruginosa* with peptides at different concentrations. Peptides are highlighted with red rectangles if inhibition zones occur around the disks.
